# Virus infection is controlled by cell type-specific sensing of murine cytomegalovirus through MyD88 and STING

**DOI:** 10.1101/2020.03.24.004788

**Authors:** Sytse J. Piersma, Jennifer Poursine-Laurent, Liping Yang, Glen N. Barber, Bijal A. Parikh, Wayne M. Yokoyama

## Abstract

Recognition of DNA viruses, such as cytomegaloviruses (CMVs), through pattern-recognition receptor (PRR) pathways involving MyD88 or STING constitute a first-line defense against infections mainly through production of type I interferon (IFN-I). However, the role of these pathways in different tissues is incompletely understood, an issue particularly relevant to the CMVs which have broad tissue tropisms. Herein, we investigated anti-viral effects of MyD88 and STING in distinct cell types that are infected with murine CMV (MCMV). Bone marrow chimeras revealed STING-mediated MCMV control in hematological cells, similar to MyD88. However, unlike MyD88, STING also contributed to viral control in non-hematological, stromal cells. Infected splenic stromal cells produced IFN-I in a cGAS-STING-dependent and MyD88-independent manner, while plasmacytoid dendritic cell IFN-I had inverse requirements. MCMV-induced natural killer (NK) cytotoxicity was dependent on MyD88 and STING. Thus, MyD88 and STING contribute to MCMV control in distinct cell types that initiate downstream immune responses.

## Introduction

Viral infections can be detected by specialized pattern recognition receptors, which recognize viral structures that are unique or otherwise absent in the subcellular location where they are detected. Nucleic acids from DNA-viruses can be detected in various organelles during infection. Some DNA viruses pass through endolysosomes where viral DNA can be recognized by toll-like receptors (TLRs), in particular TLR9, which signals through MyD88 and induces a type I interferon (IFN-I) response (Motwani, Pesiridis, & Fitzgerald, 2019). In the cytosol, infection results in exposure of viral DNA that can be recognized by cytosolic DNA sensors including cyclic GMP-AMP synthase (cGAS) and absent in melanoma 2 (AIM2) inflammasome (Tan, Sun, Chen, & Chen, 2018). cGAS signals through STING and initiates an IFN-I response, whereas AIM2 activates caspase I and instigates an IL-1β and IL-18 response. The TLRs and AIM2 pathways are primarily active in specific immune cell types. In contrast, the STING-cGAS pathway appear to be active in a broader range but not all cell types (Motwani et al., 2019). Yet, it is unclear how activation of these pathways in different cell types contributes to viral control.

Upon recognition of viral structures, IFN-I plays a central role in protection against acute infection. IFN-I mediates its anti-viral effects through stimulation of the interferon receptor, comprising of IFNAR1 and IFNAR2, and downstream STAT molecules. The resulting IFN-stimulated genes (ISGs) induce an anti-viral state, affecting cell survival and viral replication (Gonzalez-Navajas, Lee, David, & Raz, 2012; McNab, Mayer-Barber, Sher, Wack, & O’Garra, 2015). In addition, IFN-I is critical for orchestrating the subsequent innate and adaptive immune responses, through modulation of cell attraction, activation, and priming. Although human deficiencies in the IFN-I pathway are very rare, evidence suggest that IFN-I could protect against viral infections in humans. Individuals with mutations in *IFNAR2* and *STAT2* have relatively mild symptoms after infection, even though they can develop severe illness in response to live vaccines (Duncan et al., 2015; Hambleton et al., 2013). However, these deficiencies likely do not completely nullify IFN-I effects because IFNβ can signal through IFNAR1 without requiring IFNAR2 and IFN-I can signal through STAT2-independent pathways (de Weerd et al., 2013; Gonzalez-Navajas et al., 2012). Thus, it remains unclear how these different IFN-I pathways contribute to control of viral infections.

In this regard, studies of infections with the beta-herpesvirus cytomegalovirus (CMV), have been informative. Infection with human CMV (HCMV) is nearly ubiquitous worldwide (Cannon, Schmid, & Hyde, 2010). HCMV is controlled and establishes latency in healthy individuals, but HCMV can cause life-threatening disease in immunocompromised patients (Griffiths, Baraniak, & Reeves, 2015). Despite a broad tropism that allows CMV to infect a wide range of cell types, CMV is highly species-specific (Krmpotic, Bubic, Polic, Lucin, & Jonjic, 2003; Sinzger, Digel, & Jahn, 2008). Murine CMV (MCMV) in particular shares key features with HCMV and has been instructive for dissecting cytomegalovirus pathogenesis (Krmpotic et al., 2003; Picarda & Benedict, 2018). Indeed, a recent case study described a patient with deficiencies in both *IFNAR1* and *IFNGR2* who presented with bacteremia and CMV viremia (Hoyos-Bachiloglu et al., 2017). Consistent with these findings, mice deficient in *Ifnar1* and *Ifngr1* are highly susceptible to MCMV (Gil et al., 2001). *Ifnar1* deficiency in isolation resulted in a 100-fold increased MCMV susceptibility whereas *Ifngr1* deficiency did not, indicating that IFN-I plays a dominant role in controlling acute CMV infections. IFN-I production during acute MCMV infection is biphasic; initial IFN-I production peaks at 8 hours post infection (p.i.) with a second peak at 36-48 hours p.i. (Delale et al., 2005; Schneider et al., 2008). STING has been implicated in the initial IFN-I response. STING-deficient mice have decreased systemic IFNβ at 12 hours p.i. and 5-fold increased viral load at 36 hours p.i. (Lio et al., 2016). A recent study implicated Kupffer cells to be the main source for IFNβ in the liver 4 hours p.i. (Tegtmeyer et al., 2019). Besides the aforementioned immune cells, stromal cells are thought to be a major source for IFN-I in the spleen at 8 hours p.i. (Schneider et al., 2008). By contrast, MyD88-dependent pathways have been implicated in IFN-I production during the second wave (Delale et al., 2005; Krug et al., 2004). IFN-I production by plasmacytoid dendritic cells (pDCs) is dependent on TLR7 and TLR9 (Krug et al., 2004); (Hokeness-Antonelli, Crane, Dragoi, Chu, & Salazar-Mather, 2007; Zucchini et al., 2008). Consistent with the role of pDCs in IFN-I production, MyD88 is required in the hematological compartment in bone marrow chimeras (Puttur et al., 2016). However, it has been unknown which sensing pathway is responsible for IFN-I induction in the stroma and what the contribution is of the distinct sensing pathways to control MCMV infection in different tissues.

Besides its direct anti-viral effects, IFN-I is crucial for optimal NK cell function during viral infection (Orange & Biron, 1996). NK cells play a critical role in controlling MCMV infection in C57BL/6 mice, which is dependent on interactions between the Ly49H NK cell activation receptor and its MCMV-encoded ligand m157 (Arase, Mocarski, Campbell, Hill, & Lanier, 2002; Brown et al., 2001; Smith et al., 2002). However, this interaction is not sufficient to allow NK cell control of MCMV infection. IL-12 and IFN-I produced early during MCMV infection induce granzyme B and perforin protein expression in NK cells (Fehniger et al., 2007; Nguyen et al., 2002; Parikh et al., 2015), which allows them to efficiently kill virus-infected cells upon recognition of m157 through Ly49H (Parikh et al., 2015). IL-12 and IFN-I also induce IFNγ transcription, which is required for activation receptor-dependent IFNγ production (Piersma, Pak-Wittel, Lin, Plougastel-Douglas, & Yokoyama, 2019). In the absence of MyD88, Ly49H^+^ NK cells can compensate for suboptimal IFN-I production (Cocita et al., 2015), suggesting that low levels of IFN-I can still enhance NK-mediated control of MCMV. Which MCMV-sensing pathway contributes to the NK cell response is still unclear.

In the current study, we analyzed survival, viral titers, IFN-I production and NK cell responses in mice deficient in MyD88, STING or both. We also determined the contribution of both signaling pathways in different tissues to their anti-viral effects, and elucidated a role for cGAS in these responses.

## Results

### MyD88 and STING-dependent pathways control MCMV infection in vivo

We set out to investigate the contribution of STING- and MyD88-dependent pathways in controlling MCMV infection by analyzing the morbidity and mortality in wildtype C57BL/6 (WT), MyD88-deficient (MyD88 KO), and STING-deficient (STING GT) mice as well as mice deficient in both MyD88 and STING (DKO) that were infected with 50,000 PFU MCMV (Figure 1). Consistent with previously published data (Lio et al., 2016), WT mice lost approximately 10% of weight by 3 days p.i. after which they recovered (Figure 1A). Here we observed that STING GT mice showed more pronounced weight loss compared to WT mice, but were also able to recover. MyD88 KO mice showed delayed weight loss as compared to WT mice, indicating that the initial weight loss in WT mice was caused by immunopathology mediated by MyD88. The weight curves of DKO mice overlapped with MyD88 KO mice, suggesting that the extra weight loss in the STING deficient mice as compared to WT mice is likely MyD88-dependent. Both WT and STING GT mice were able to control and survive viral infection upon challenge with MCMV (Figure 1B). MyD88 KO mice were moderately resistant to the infection as 37% of the mice died between days 6 and 7. In contrast, the majority (70%) of DKO mice succumbed to the infection. These data show that both STING and MyD88 significantly contribute to control of MCMV infection *in vivo.*

**Figure 1:**
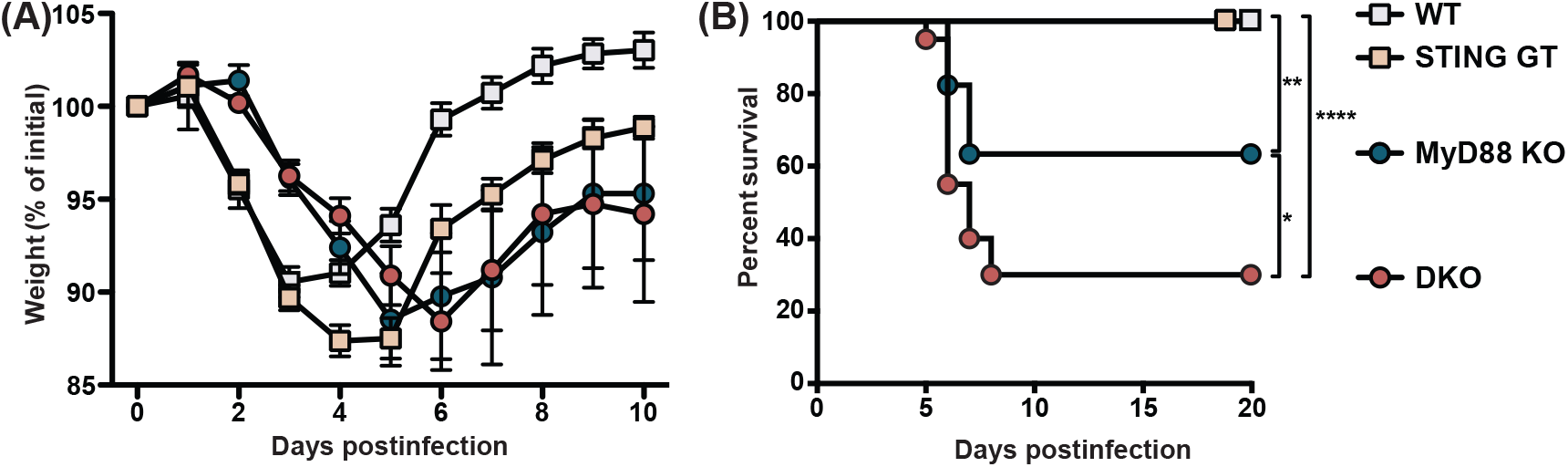
MyD88 and STING control morbidity and mortality during MCMV infection. Mice were infected with 50,000 PFU MCMV WT-1, weight loss and survival was monitored over time. (**A**) Weight loss over time in wildtype (n=12), STING-deficient (STING GT, n=21), MyD88-deficient (MyD88 KO, n=9) and mice deficient in both STING and MyD88 (DKO; n=14). (**B**) Survival curves of wildtype (n=17), STING GT (n=18), MyD88 KO (n=17) and DKO mice (n=20). Cumulative data of 3 independent experiments. Error bars indicate SEM; *p<0.05, **p<0.01, ****p<0.0001.

### STING contributes in both the hematological and radio-resistant compartments in controlling viral load

To investigate the contribution of STING and MyD88 in different organs we analyzed viral loads in the spleen and liver, the initial organs of replication after infection (Hsu, Pratt, Akers, Achilefu, & Yokoyama, 2009; Sacher et al., 2008). In the spleen we observed a modest but significant increase (6.9-fold) in viral load in MyD88 KO mice two days p.i., whereas the spleens of DKO mice contained 84-fold higher viral copies compared to WT controls (Figure 2A). By 5 days p.i. we observed an 85-fold increase in viral load in the spleens of MyD88-deficient and 1901-fold increase in viral load in DKO, both as compared to WT controls (Figure 2B). We did not observe significant differences in STING-deficient animals, but we observed a 23-fold increase in viral load in DKO spleens compared to MyD88 KO, indicating that STING contributes to viral control in the absence of MyD88. In the liver, we were unable to detect significant differences in viral load at 2 days p.i. (Figure 2A). By day 5, we observed a 221-fold increase in DKO and 51-fold increase in MyD88 KO viral load compared to WT controls (Figure 2B). Taken together, these data indicate that the STING and MyD88 pathways contribute to viral control at early timepoints, particularly in the spleen and to a lesser extent in the liver.

**Figure 2:**
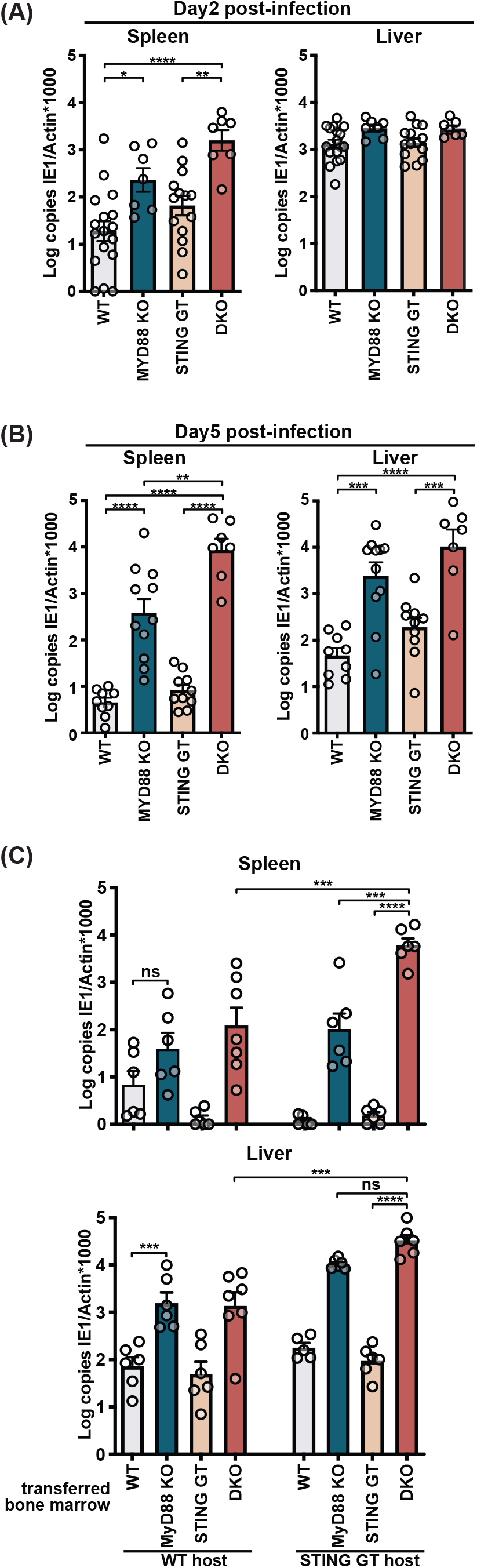
STING contributes to control of MCMV in the hematological and stromal compartment, whereas MyD88 in the hematological compartment potently controls infection. Mice were infected with 50,000 PFU (**A**) and (**B**) or 20,000 PFU (**C**) MCMV. Viral load was quantified 2 days (**A**) or 5 days (**B**) and (**C**) p.i. (**C**) Indicated bone marrow was adoptively transferred into irradiated wildtype (WT) or STING-deficient (STING GT) hosts. Bone marrow chimeras were infected 6 weeks post transfer and viral load was analyzed 5 days p.i. Each panel shows cumulative data of 2 independent experiments. Error bars indicate SEM; ns, not significant, *p<0.05, **p<0.01, ***p<0.001, ****p<0.0001.

MyD88 has been reported to be required in the hematological compartment, but not in the radio-resistant compartment (Puttur et al., 2016), yet it is unclear which compartment(s) requires STING-dependent pathways. To investigate the contributions of STING and MyD88 dependent pathways in these compartments, we generated bone marrow (BM) chimeras of either or both knockout BM into irradiated WT or STING GT hosts and analyzed viral load day 5 p.i. (Figure 2C). While WT hosts reconstituted with WT BM controlled viral load similar to WT control mice (Figure 2C vs Figure 2 - Figure supplement 1A), reconstitution of WT hosts with MyD88-deficient BM resulted in elevated viral loads compared to reconstitution with WT BM (Figure 2C), consistent with previous published results (Puttur et al., 2016). We also observed that the contribution of MyD88 in the hematological compartment was particularly overt in the absence of STING, revealed by comparison of DKO BM into STING GT host versus STING GT BM into STING GT host, which resulted in a 3882-fold difference in the spleen and 344-fold in the liver, respectively. STING also had anti-viral effects in the hematological compartment, evident by comparing DKO BM into STING GT host versus MyD88 KO BM into STING GT host, which revealed a 59-fold difference in the spleen. STING played also a role in the radio-resistant compartment in both spleen and liver, revealed by comparison of DKO BM into WT host versus DKO BM into STING GT host, for which we observed a 49-fold and 24-fold differences in the spleen and liver, respectively. Jointly, the BM chimeras reveal an evident role for MyD88 in the hematological compartment, while STING contributes to viral control in both the hematological and radio-resistant compartments, most explicitly in the spleen.

### Multiple cell populations produce IFN-I in response upon MCMV infection

Type I IFNs are induced in response to triggering of pathogen recognition receptors that signal through MyD88 and STING and are key players in the initial anti-viral response. To investigate the IFN-I response in virus-infected cells we made use of a reporter virus that expresses GFP under the IE1 promoter (Henry et al., 2000). We analyzed initial times (8- and 36-hours p.i.) and focused on stromal cell and dendritic cell (DC) populations, which are the major cell types infected at these timepoints (Hsu et al., 2009). Consistent with previous published data, we detected infection of the stromal cell but not DC compartment at 8-hours p.i. (Figure 3A). At 36-hours p.i., the percentage of infected stromal cells increased substantially and infected DCs were detected as well. Infected DCs included conventional DC (cDC; CD11c high in Figure 3A) and plasmacytoid DC (pDC; CD11c low in Figure 3A) populations. Based on these data, we sorted and analyzed infected and uninfected populations at 36-hours p.i. for IFNα and IFNβ transcripts by quantitative PCR (Figure 3 – Figure supplement 1). The infected stromal cells (GFP^+^) specifically expressed *Ifna* and *Ifnb1* transcripts, which were not detected in the uninfected (GFP^-^) cells (Figure 3B). Infected DCs also expressed transcripts for *Ifna* and *Ifnb1* but *Ifna* transcripts were also detected in GFP^-^ DCs isolated from MCMV-infected animals (Figure 3C). These data suggest that *Ifna* transcripts in uninfected DCs are produced (in part) as a feedforward loop (McNab et al., 2015), whereas *Ifnb1* transcripts are specifically produced in response to viral detection in infected DCs as well as stromal cells. Based on these data we chose to investigate the role of STING and MyD88 on IFNβ production by different cell types.

**Figure 3:**
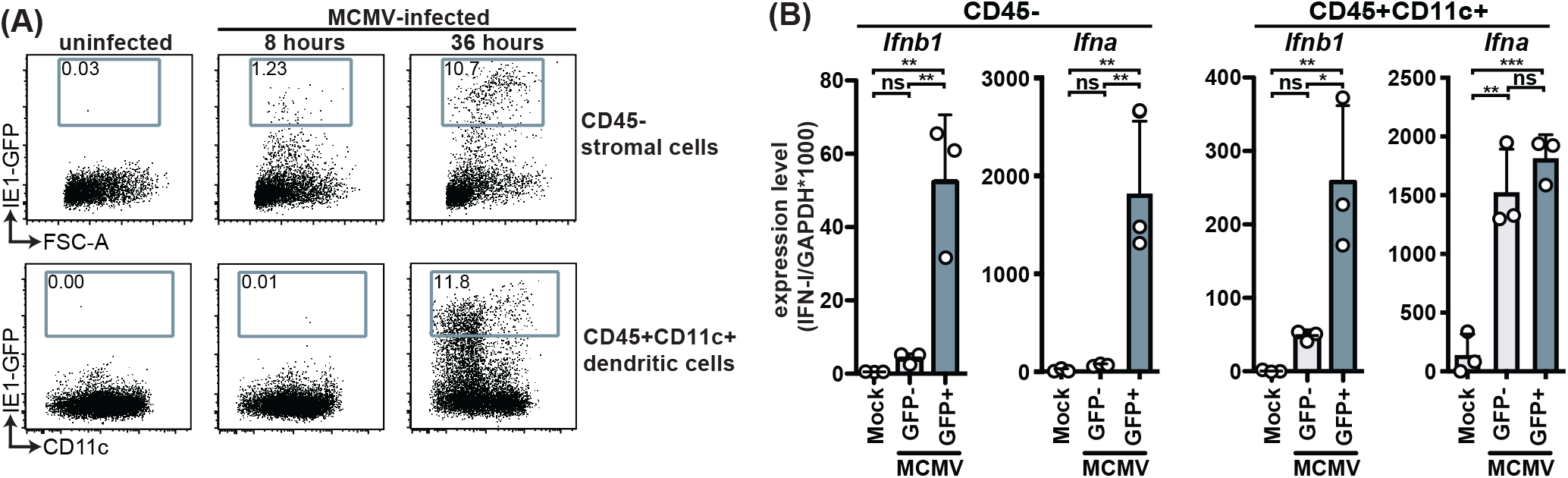
MCMV-infected cells specifically produce IFNβ upon infection. WT mice were infected with 100,000 PFU MCMV IE1-GFP reporter virus. (**A**) Analysis of GFP expression in CD45^-^ stromal cells and CD45^+^CD11c^+^ DC at 8 hours and 36 hours p.i. (**B**) GFP^+^ and GFP^-^ stromal cells and DC were FACS-sorted 36 hours p.i. and *Ifnb1* and pan- *Ifna* transcript levels were quantified by real-time PCR. Both panels show representative experiments from 2 independent experiments. Error bars indicate SD; ns, not significant, *p<0.05, **p<0.01, ***p<0.001.

### IFNβ is produced by pDCs in a MyD88-dependent but STING-independent manner during infection

To evaluate the role of STING and MyD88 on IFN-I production by individual cells upon infection, we backcrossed MyD88 KO, STING GT, and DKO mice to the IFNβ-YFP reporter mouse (Scheu, Dresing, & Locksley, 2008). Approximately 20% of the pDCs were YFP^+^, indicating at least this percentage of infected pDCs produced IFNβ in response to MCMV infection, whereas much fewer cDCs produced IFNβ because less than 1% of cDCs were YFP^+^ (Figure 4B). Consistent with previous studies of primary pDC *in vitro* (Krug et al., 2004), we observed that the production of IFNβ by pDCs was solely dependent on MyD88 because MyD88 KO mice were unable to induce detectable YFP (IFNβ) in pDCs. By contrast, here we found that STING GT mice did not significantly affect pDC IFNβ production, indicating that MyD88 functions in these cells without requiring STING-dependent pathways. On the other hand, both STING and MyD88 seemed to affect IFNβ reporter levels in the few YFP^+^ cDCs, although the differences were not significant (Figure 4B). Nonetheless, these results indicate that MyD88-dependent sensing of MCMV dictated the IFNβ response in pDCs, but it remained unclear how MyD88- and STING-dependent pathways contribute to IFNβ production in stromal cells.

**Figure 4:**
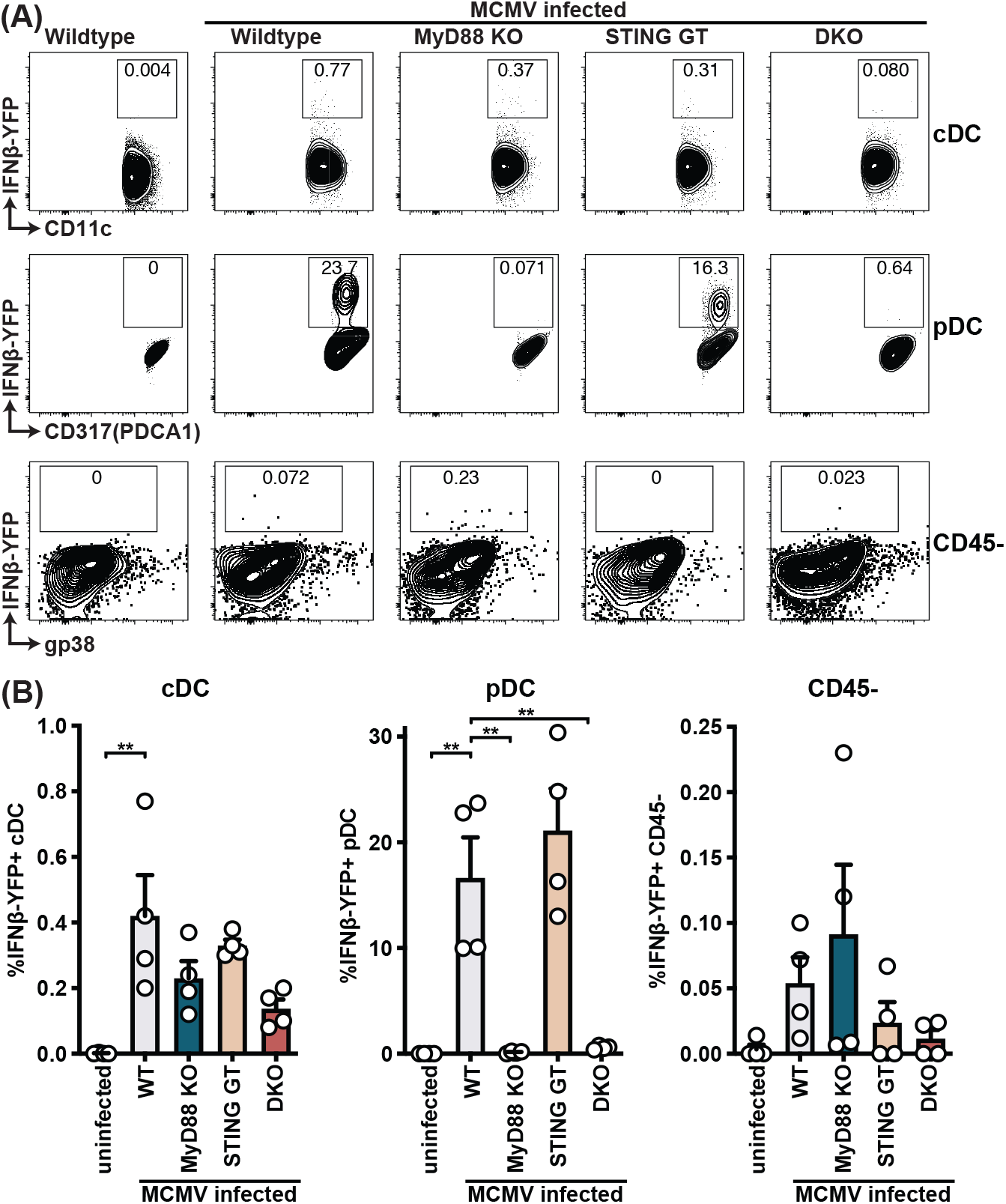
pDCs produce IFNβ in a MyD88-dependent but STING-independent manner in IFNβ-YFP reporter mice. IFNβ-YFP reporter mice were backcrossed to MyD88- (MyD88 KO), STING- (STING GT) and double-deficient (DKO) mice. Animals were infected with 200,000 PFU WT1 MCMV and analyzed 48 hours post infection. Spleens were digested to a single cell suspension, stained and analyzed by flow cytometry. Error bars indicate SD; **p<0.01.

### IFNβ is produced by stromal cells in a STING-dependent but MyD88 independent manner during infection

Since we were unable to find YFP^+^ infected stromal cells, which might be due to a detection level issue in these cells *in vivo* (Figure 4), we turned to *in vitro* infection of primary stromal cells. To this end, we isolated primary splenic fibroblast and challenged them with MCMV at MOI 5 (Figure 5A). Indeed, splenic fibroblasts readily expressed 8000-fold increase in *Ifnb1* transcripts by qPCR at 8 hours p.i. To determine the role of key innate sensing components, we turned to mouse embryonic fibroblasts (MEFs) that were genetically deficient in these components. Consistent with primary splenic fibroblasts, MEF expressed *Ifnb1* transcripts upon MCMV infection (Figure 5B), reaching levels to those detected in primary splenic fibroblasts. We further observed that *Ifnb1* expression was independent of MyD88 and TRIF, indicating that TLRs do not contribute to IFNβ production in fibroblasts even though *Ifnb1* expression was dependent on IRF3/7 and TBK1, which is consistent with cytosolic sensing of MCMV. Using MEF lines with 2 different mutations in STING (Ishikawa & Barber, 2008; Sauer et al., 2011), we found that the IFNB1 response was instead dependent on STING. However, IFNB1 production was independent of MAVS (also known as CARDIF and IPS-1), suggesting that the IFN-response is independent of the cytosolic RNA sensors (Tan et al., 2018). Finally, we investigated the role of cytosolic DNA sensors and found that fibroblast sensing of MCMV was dependent on cGAS, but independent of ZBP1 and DNA-PK. To confirm that the cGAS pathway also is involved in adult splenic stromal cells, we analyzed *Ifnb1* expression in cGAS-deficient primary splenic fibroblasts (Figure 5C). Indeed, cGAS-deficient splenic fibroblasts were unable to express *Ifnb1* in response to MCMV challenge, indicating that the STING-cGAS-dependent pathway is responsible for the IFNβ response in the stromal cell compartment. To validate that these pathways are also involved in IFNβ protein production and secretion, we analyzed cell culture supernatants at 48 hours p.i. with MCMV MOI 0.5 (Figure 5E). WT MEF secreted IFNβ in response to MCMV infection, but neither STING nor cGAS-deficient MEFs produced IFNβ. Collectively, these results strongly suggest that the stromal cell compartment produces IFNβ in a STING-cGAS dependent but MyD88-independent manner.

**Figure 5:**
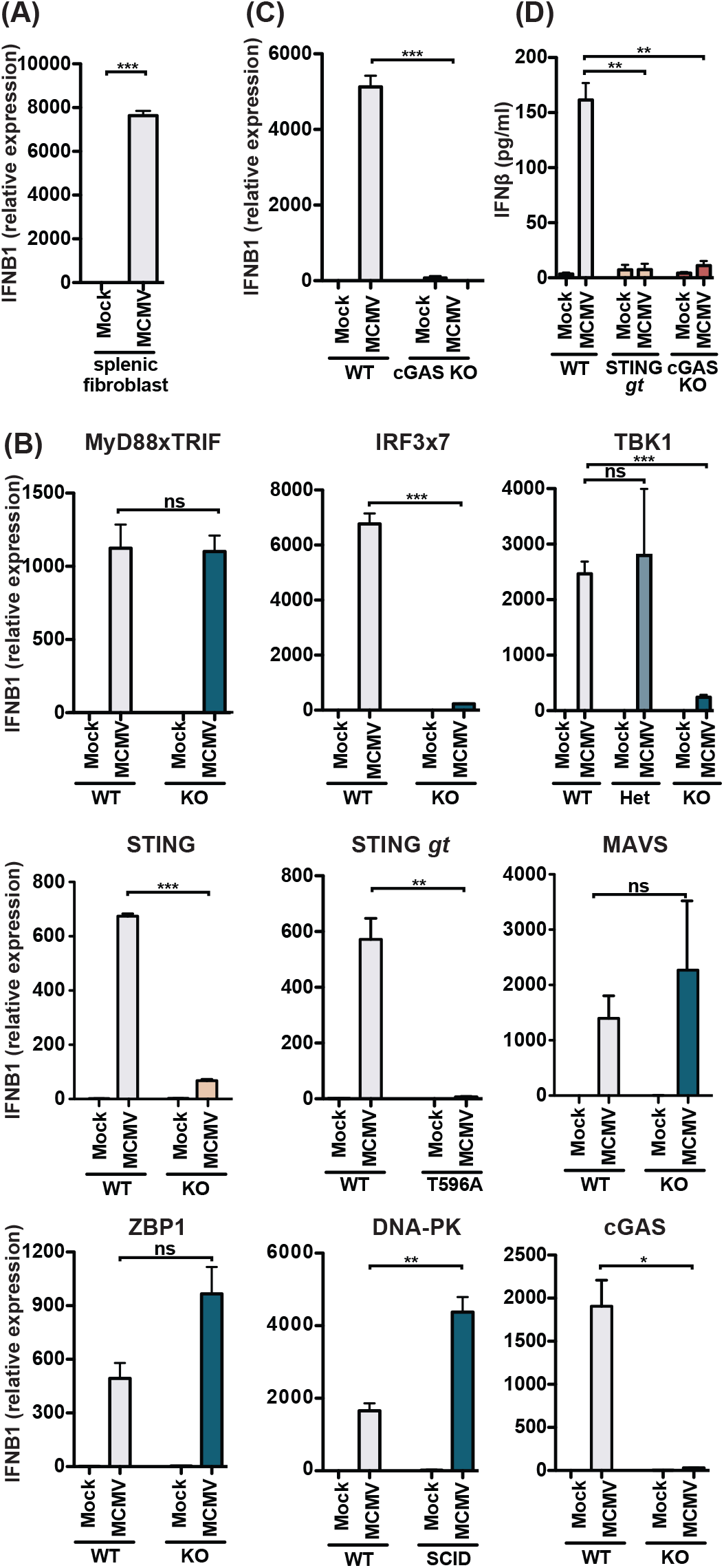
MCMV-induced fibroblast IFNβ is triggered by cGAS-STING-dependent but MyD88-Trif-MAVS-independent mechanisms. (**A**) IFNB1 mRNA levels of primary splenic fibroblasts infected with WT1 MCMV (MOI=5) 8 hours post-infection. (**B**) IFNB1 mRNA levels of murine embryonic fibroblasts (MEF) from wildtype (WT) or indicated deficient mice were infected and analyzed as in (A). (**C**) IFNB1 mRNA levels in infected WT or cGAS-deficient primary splenic fibroblasts, analyzed as in (A). (**D**) Secreted IFNβ by WT or indicated gene deficient MEF, infected with MCMV (MOI=0.5); supernatant was analyzed 48 hours p.i. by ELISA. Panels show representative experiments from 2 independent experiments performed in duplicate. WT, STING GT, and TBK1-, MAVS-, ZBP1-, DNA-PK-, and cGAS-deficient MEF represent data from 2 independent MEF preparations. Error bars indicate SEM; ns, not significant, *p<0.05, **p<0.01, ***p<0.001.

### MyD88 and STING contribute to NK cell cytolytic potential

We previously reported that both IFN-I and IL-12 act directly on NK cells to induce perforin (Prf) and granzyme B (GzB) protein levels, thereby increasing NK cell cytolytic potential, which was required for Ly49H-dependent control of MCMV infection (Parikh et al., 2015). Moreover, IL-12 production in response to MCMV has been reported to be dependent on MyD88 (Krug et al., 2004), and thus contributed to the phenotypes observed in MyD88 KO mice independent of IFN-I. Here we investigated the role of MyD88 and STING in increasing NK cell reactivity during MCMV infection. Consistent with previous reports (Fehniger et al., 2007; Orange, Wang, Terhorst, & Biron, 1995; Parikh et al., 2015), we observed increased levels of NK cell GzB, Prf and IFNγ at 48 hours p.i. (Figure 6A). At this time point, NK cell production of IFNγ is dependent on IL-18, which signals through MyD88 (Adachi et al., 1998; Pien, Satoskar, Takeda, Akira, & Biron, 2000). Indeed, NK cell IFNγ production was dependent on MyD88, whereas STING did not impair IFNγ production, and rather increased the IFNγ response (Figure 6B). This potentially could be due to a relatively small increase in viral load at these timepoints. Expression of both GzB and Prf followed a similar pattern, as the vast majority of increased expression was dependent on MyD88, whereas STING did not overtly contribute to the production of these lytic proteins (Figure 6B). Finally, we analyzed NK cell cytolytic capacity using a 3-hour *in vivo* target cell rejection assay. We previously reported that MCMV increased m157-target cell rejection in an IL-12- and IFN-I-dependent manner (Parikh et al., 2015). Consistent with our previous data, m157-specific target cell rejection increased 3 days post-MCMV infection from 50% to 80% (Figure 6C). MHC-I-deficient cell (“missing self”) rejection was higher and increased from 30% to 90%. MyD88 KO or STING GT mice did not display substantial differences in target cell rejection, but DKO mice substantially decreased NK cell cytolytic capacity with m157-specific rejection showing levels of uninfected mice. Similarly, MHC-I specific rejection was decreased in double versus single deficient mice. Together, these data indicate that both MyD88 and STING-dependent pathways contribute to NK cell cytolytic potential, albeit that MyD88 predominantly affects production of Prf and GzB.

**Figure 6:**
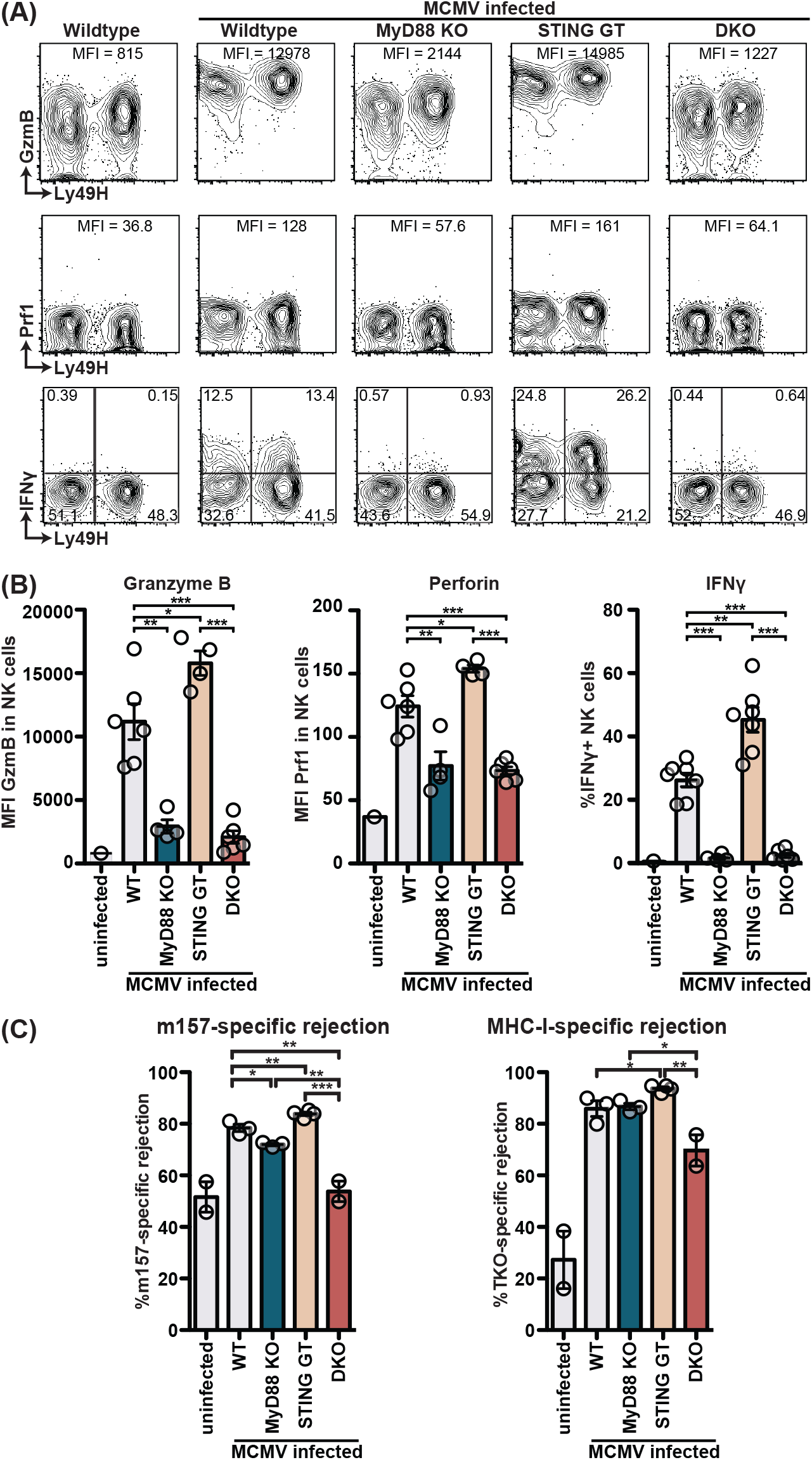
MyD88 and STING are required for NK cell cytolytic capacity during MCMV infection. (**A** and **B**) Mice deficient in MyD88 and/or STING were infected with MCMV and 2 days later splenocytes were harvested and analyzed for GzmB, Prf1, and IFNγ expression by FACS. Representative contour plots of individual mice are shown in (**A**) and quantification for multiple mice is shown in (**B**). (**C**) Differentially labelled WT, m157-Tg and MHC-I deficient splenocytes were adoptively transferred into indicated day 3-infected mice. Specific rejection was analyzed 3 hours post-transfer in the spleen. Representative experiments from 2 independent experiments per panel are shown. Error bars indicate SEM; ns, not significant, *p<0.05, **p<0.01, ***p<0.001.

## Discussion

Herein we describe that MCMV infection can be sensed by both STING and MyD88-dependent pathways which contribute to viral control in response to lethal challenge. While we confirmed the strong role of MyD88 in the hematological compartment, especially in splenic pDCs, we found that STING contributes in both the hematological and the previously unappreciated stromal cell compartment. Using primary splenic stromal cell cultures, we identified a role for cGAS-STING-dependent, but MyD88-independent IFN-I production in response to MCMV infection. Finally, we found that both MyD88 and STING-dependent pathways contribute to increased NK cell cytolytic function during infection. Thus, our findings indicate that cytomegalovirus infection is sensed by distinct sensing pathway depending on the infected cell type and that these pathways constitute a multi-layer antiviral defense.

Cytomegaloviruses have a broad tropism and a broad range of infected cell types have the capacity to produce IFN-I in response to infection. However, IFN-I production has been most well characterized in myeloid cells, including pDCs and Kupffer cells. IFN-I production by pDCs upon MCMV infection *in vitro* is dependent on TLR9 and MyD88 (Krug et al., 2004). Using IFNβ reporter mice, we were able to confirm that pDCs were the major source of IFNβ in the spleen and that this was dependent on MyD88. Furthermore, we observed that this IFN-I production was independent of STING. Early after infection, Kupffer cells in the liver produce IFN-I in a STING-dependent, but TLR-independent manner (Tegtmeyer et al., 2019). Hepatocytes are a major target for infection by MCMV(Sacher et al., 2008), yet they do not induce detectable levels of IFNβ (Tegtmeyer et al., 2019). However, hepatocytes do not express STING (Thomsen et al., 2016), likely explaining the lack of IFNβ production in hepatocytes in response to MCMV infection. In our bone marrow chimeras, we observed a role for STING in the radioresistant compartment in the liver. Hepatic stromal cells, including endothelial cells, are infected with MCMV (Sacher et al., 2008), providing likely contributors, apart from hepatocytes per se, to control MCMV in a STING-dependent manner in the liver.

We observed a stronger effect of STING in the splenic stromal cell compartment compared to the liver stromal cell compartment. IFN-I produced by splenic stromal cells have previously been reported to be dependent on lymphotoxin (LT) β expression by B cells, independent of TLR signaling (Sacher et al., 2008). Consistent with these findings, our findings revealed that stromal cell IFN-I is cGAS-STING-dependent. Besides the anti-viral role for STING in the stromal cell compartment, we observed that IFN-I production by infected primary splenic stromal cells was cGAS-STING-dependent. The primary stromal cells did not require interactions with B cells to produce IFN-I *in vitro*. However, it remains to be determined how LT intersects with the STING pathway *in vivo*, particularly since LT has been reported to be required for cell survival during MCMV infection (Banks et al., 2005), potentially providing a window where infected stromal cells survive long enough to produce IFN-I. Additional studies are required to further define these experimentally complex interactions.

Herpesviruses dedicate a large part of their genome to immune evasion strategies, including strategies that act on cellular immunity and intrinsic cellular defenses (Powers, DeFilippis, Malouli, & Fruh, 2008). MCMV has been reported to interfere with the DNA sensing pathway at different steps, m152 binds to STING and interferes with its trafficking (Powers et al., 2008), whereas m35 targets NFκB-mediated transcription (Chan et al., 2017). Deletion of these ORFs individually in MCMV results in stronger IFN-I responses upon infection *in vivo.* Infection with MCMV deleted in both ORFs potentially facilitates visualization of the IFN-I production by the cell types under study. Despite these immune evasion strategies, WT MCMV induces an IFN-I response that is potent enough to control virus infection, therefore we chose to use WT MCMV in the current study.

Infection with lethal dose of MCMV resulted in about a third of the MyD88-deficient mice succumbing to infection, which is consistent with previously published results (Delale et al., 2005). However, a recent study did not observe a lethality phenotype in mice deficient in MyD88 and TRIF, unless mice also lacked MAVS (Tegtmeyer et al., 2019). The latter study utilized tissue culture-derived MCMV in contrast to salivary gland extracted MCMV in the former and our study. Additionally, Tegtmeyer et al. used a mutant MCMV that lacked m157, whereas we used m157-sufficient virus for infections monitoring survival. Since IFN-I impacts NK cell-dependent MCMV-control via m157 recognition (Parikh et al., 2015), the use of WT MCMV allowed us to evaluate the effect of the virus-sensing pathways on NK cell function.

IFN-I and IL-12 produced in response to MCMV infection are required for full NK cell cytolytic capacity, through induction of GzB and Prf (Parikh et al., 2015). Consistent with previous reports (Krug et al., 2004; Puttur et al., 2016), we found that MyD88-deficient mice expressed low levels of NK cell GzB, Prf and IFNγ in response to MCMV infection but NK cytolytic potential *in vivo* was not substantially affected. However, MCMV-infected mice deficient in both STING and MyD88 displayed reduced NK cell cytolytic activity against m157-expressing and MHC-I-deficient target splenocytes (Cocita et al., 2015). Thus, MyD88- and STING-dependent sensing of MCMV both contribute to signal to NK cells to enhance their cytolytic function in order to efficiently clear MCMV-infected target cells.

## Materials and Methods

### Mice

C57BL/6 (stock number 665) and BALB/c (028) mice were purchased from Charles River Laboratories. The following mouse strains were purchased from Jackson Laboratories: STING golden ticket (017537), IFNβ-YFP reporter mice (010818), DNA-PK SCID (001913), and β2m KO (002087). m157-Tg mice were generated and maintained in-house. H-2K^b^ KO x H-2D^b^ KO (4215) mice were purchased from Taconic Farms. MyD88 KO, TBK1 KO (nbio156), and ZBP1 KO (nbio155) mice were kindly provided by S. Akira (Osaka University, Osaka, Japan) through the JCRB Laboratory Animal Resource Bank of the National Institute of Biomedical Innovation (Adachi et al., 1998; Hemmi et al., 2004; Ishii et al., 2008) and were maintained on a C57BL/6 background. IPS1 KO mice on the C57BL/6 background were kindly provided by Michael Gale (University of Washington, Seattle, WA, USA). Mice deficient for cGAS were kindly provided by Herbert Virgin (Vir Biotechnology, San Francisco, CA, USA)(Schoggins et al., 2014). Triple MHC Class I KO mice (TKO) were generated by crossing β2m KO mice to H-2K^b^ KO x H-2D^b^ KO mice. STING GT mice were crossed to MyD88 KO to generate DKO mice. Subsequently DKO and single KO mice were crossed with IFNβ-YFP reporter to generate IFNβ-YFP on the various KO backgrounds. All mice were maintained in-house in accordance with institutional ethical guidelines. Age- and sex-matched mice were used in all experiments.

### Cell lines

3T12 cells (ATCC CCL-164) were maintained in DMEM supplemented with newborn calf serum, L-glutamine, penicillin, and streptomycin and were used for production of tissue culture derived MCMV and tittering of virus stocks. All MEF were maintained in RPMI supplemented with fetal bovine serum, L-glutamine, penicillin, and streptomycin. IRF3/7 KO MEF were kindly provided by Michael S Diamond (Washington University in St Louis, MO, USA). STING KO MEF have been described before (Ishikawa, Ma, & Barber, 2009). All other MEF lines were generated from day 11.5-13.5 embryos, at least 2 independent lines were generated per genotype. To generate splenic fibroblasts, spleens were minced and digested with Liberase TL, adherent cells were cultured for 3-6 weeks to obtain pure fibroblast populations.

### In vivo virus infections

For *in vivo* studies salivary gland MCMV (sg-MCMV) of the WT-1 strain, a subcloned Smith strain (Cheng, Valentine, Gao, Pingel, & Yokoyama, 2010), was used for infections unless otherwise indicated. Where indicated, MCMV that expressed GFP under the IE1 promotor was used to visualize infected cells (Henry et al., 2000). This reporter virus contained a mutation in m157. All viral strains for *in vivo* infections were propagated in BALB/c mice; virus was isolated from salivary glands and titers were determined as previously described (Brune, Hengel, & Koszinowski, 2001; Jonjic, 2001). Mice were infected with indicated dose of MCMV intraperitoneally in 200μl PBS. For survival studies weight was monitored daily and mice were sacrificed when more than 30% of initial weight was lost, in accordance to animal protocol. Viral load analysis was performed as previously described (Parikh et al., 2015). Briefly, RNA-free organ DNA was isolated using Puregene extraction kit (Qiagen). 160 ng DNA was quantified for MCMV IE1 (Forward: 5’-CCCTCTCCTAACTCTCCCTTT-3’; Reverse: 5’-TGGTGCTCTTTTCCCGTG −3’; Probe: 5’-TCTCTTGCCCCGTCCTGAAAACC-3’; IDT DNA) and host *Actb* (Forward: 5’-AGCTCATTGTAGAAGGTGTGG-3’; Reverse: 5’ - GGTGGGAATGGGTCAGAAG-3’; Probe: 5’-TTCAGGGTCAGGATACCTCTCTTGCT-3’; IDT DNA) against plasmid standard curves using TAQman universal master mix II on a StepOnePlus real time PCR system (Thermo Fisher Scientific).

### Bone marrow chimeras

C57BL/6 and STING GT mice were irradiated with 950 rad by an x-ray irradiator and were intravenously with 5 million of the indicated genotype donor bone marrow cells. Chimeric mice were given antibiotic water (sulfamethoxazole/trimethoprim) for 4 weeks. 6 weeks post-irradiation mice were infected with MCMV and analyzed for viral load at 5 days p.i. We observed greater sensitivity of reconstituted BM chimeric mice to infections than mice not subjected to the BM transplant procedure in our facility so we infected reconstituted mice with a lower dose of MCMV (20,000 PFU) as compared to non-chimeric mice.

### In vitro virus infections

For *in vitro* studies, pelleted tissue culture-derived MCMV was prepared and viral titers were determined as previously described (Brune et al., 2001). 200,000 cells were plated in a 6-well plate overnight and were infected with 200μl of MCMV at MOI 5 for RNA analysis and MOI 0.5 for supernatant analysis for 1 hour, after which wells were washed with PBS to remove free virus and 2 ml fresh culture media was added. Cells were lysed in the wells with 1 ml trizol after an additional 5 hours culture for RNA analysis. Samples were stored at −80°C until analysis. Supernatants were harvested 48 hours after culture and analyzed for IFNβ by ELISA (Biolegend) according to manufacturer protocol.

### Flow Cytometry

Fluorescent-labeled antibodies used were anti-NK1.1 (clone PK136), anti-NKp46 (29A1.4), anti-CD3 (145-2C11), anti-CD19 (eBio1D3), anti-CD31 (390), anti-PDCA1 (eBio129c), anti-gp38 (eBio8.1.1), anti-CD45 (30-F11), anti CD11c (N418), anti-Ly49H (3D10), anti-Perforin (eBioOMAK-D), anti-Granzyme B (GB12), and anti-IFNγ (XMG1.2), all from Thermo Fisher Scientific. For analysis of splenic dendritic and stromal cells, spleens were digested with 1 mg/ml Liberase TL and DNAse -I (Millipore Sigma) for 45 minutes with mechanical dissociation with a pipette every 15 minutes to obtain a single cell suspension. For analysis of NK cells, spleens were crushed through a 70μm cell strainer to obtain a single cell suspension. Red blood cells (RBC) in all samples were lysed with RBC lysis buffer. Cells for analysis were first stained with fixable viability day (Thermo Fisher Scientific). Subsequently, cell surface molecules were stained in 2.4G2 hybridoma supernatant to block Fc receptors. For intracellular staining, cells were fixed and stained intracellularly using the Cytofix/Cytoperm kit (BD Biosciences) according to manufacturer’s instructions. Samples were acquired using FACSCanto (BD Biosciences) and analyzed using FlowJo software (Treestar). NK cells were defined as Viability-NK1.1^+^CD3^-^CD19^-^. Where indicated, cells were sorted on a FACSaria (BD Biosciences) into media and subsequently lysed in Trizol for RNA analysis.

### RNA analysis

RNA was isolated from cultured or sorted cells using Trizol according to manufacturer instruction (Thermo Fisher Scientific). Contaminating DNA was removed using Turbo DNAse, and cDNA was synthesized using Superscript III using oligo(dT) (Thermo Fisher Scientific). Quantification was performed for *Ifnb1* (Mm00439546_s1; Thermo Fisher Scientific), *(pan)Ifna* (Forward: 5’-CTTCCACAGGATCACTGTGTACCT-3’; Reverse: 5’-TTCTGCTC TGACCACCTCCC-3’; Probe: 5’-AGAGAGAAGAAACACAGCCC CTGTGCC-3’; IDT DNA)(Samuel & Diamond, 2005) and *Gapdh* (Mm99999915_g1; Thermo Fisher Scientific) against plasmid or pooled standard curves using TAQman universal master mix II on a StepOnePlus real time PCR system (Thermo Fisher Scientific).

### In vivo cytotoxicity assay

Target splenocytes were isolated from C57BL/6, m157-Tg, and MHC-I deficient (TKO) mice and differentially labelled with CFSE, CellTrace violet, and CellTrace far red (Thermo Fisher Scientific). Target cells were mixed at a 1:1:1 ratio and 3×10^6^ target cells were injected i.v. into naïve or day 3 MCMV-infected mice. 3 hours after challenge splenocytes were harvested and stained. The ratio of target (m157-tg or TKO) to control (C57BL/6) viable CD19^+^ cells was determined by flow cytometry. Target cell rejection was calculated using the formula [(1-(Ratio(target:control)_sample_/Ratio(target:control)_NK depleted_)) × 100]. Average of two NK1.1-depleted mice served as control.

### Statistical analysis

Statistical analysis was performed using Prism (GraphPad software). Survival curves were compared using Log-Rank (Mantel-Cox) tests, other comparisons were performed using one-way ANOVA with Bonferroni’s multiple comparisons tests to calculate P values. Error bars in figures represent the SEM. Statistical significance was indicated as follows: ****, P < 0.0001; ***, P < 0.001; **, P < 0.01; *, P < 0.05; ns, not significant.

## Acknowledgements

We thank Beatrice Plougastel-Douglas for critically reading the manuscript.

This work was supported by NIH grant R01-AI131680 to W.M.Y. and S.J.P. was supported by the Netherlands Organisation for Scientific Research (Rubicon grant 825.11.004).

The authors declare no competing financial interests. S.J.P., B.A.P., G.N.B., and W.M.Y. designed the research. S.J.P., J.L., L.Y. performed experiments. S.J.P. and W.M.Y. wrote the manuscript.

---

**Figure 2 – Figure supplement 1:**
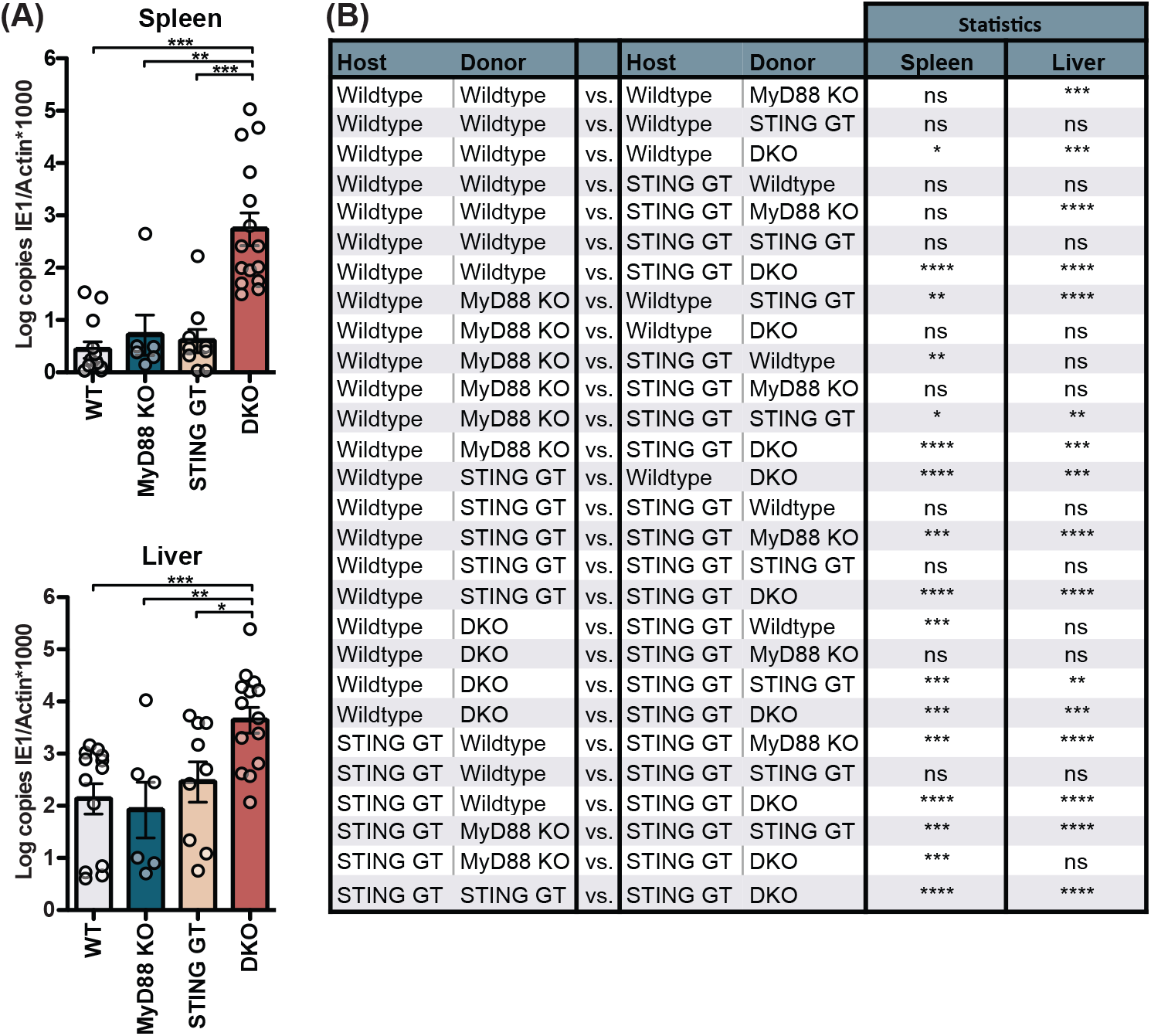
Viral load for 20,000 PFU infection 5 days p.i. and extended statistical analysis for bone marrow chimeras. (**A**) Mice were infected with 20,000 PFU and viral load was quantified 5 p.i. (**B**) Extended statistical analysis for bone marrow chimeras presented in Figure 2C. Each panel shows cumulative data of 2 independent experiments. Error bars indicate SEM; ns, not significant, *p<0.05, **p<0.01, ***p<0.001, ****p<0.0001.

**Figure 3 – Figure supplement 1:**
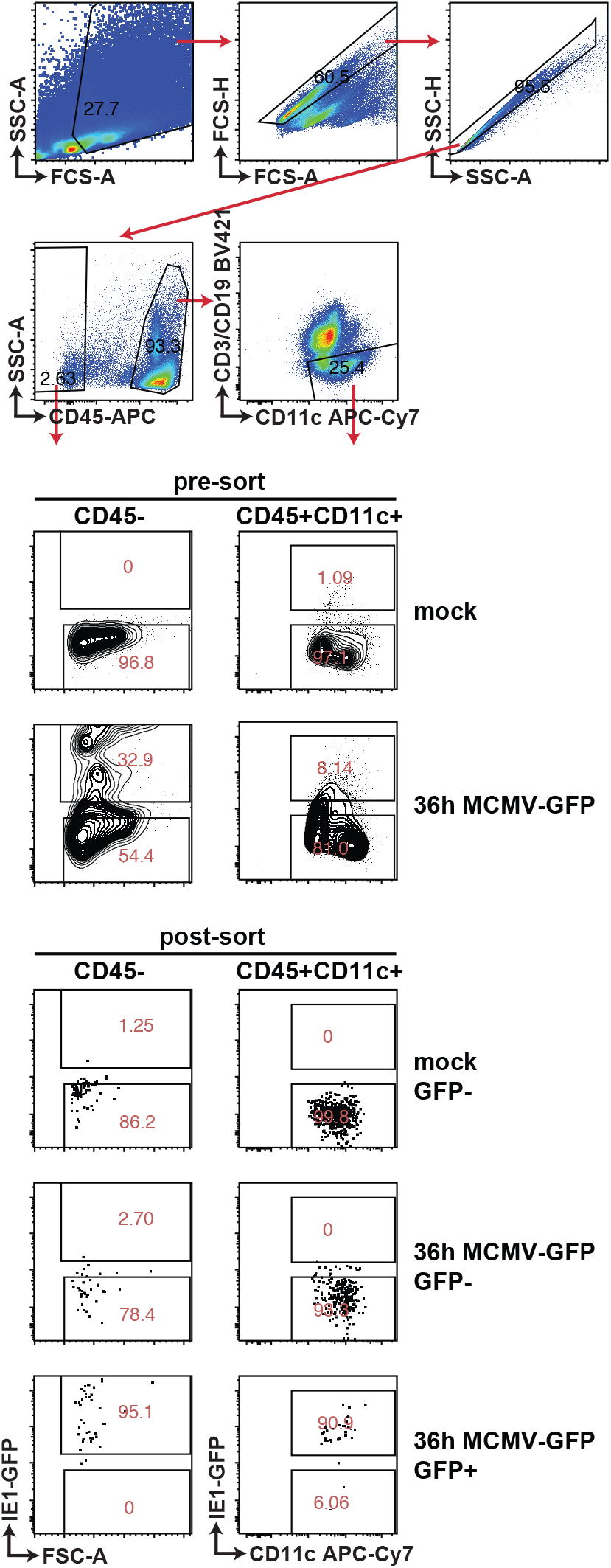
Gating strategy and purity of sorted cell populations. WT mice were infected with 100,000 PFU MCMV IE1-GFP reporter virus. GFP^+^ and GFP^-^ stromal cells and DC were FACS-sorted 36 hours p.i. Representative gating strategy and purity of sorted cells that were used in Figure 3BC are shown.

**Figure 4 – Figure supplement 1:**
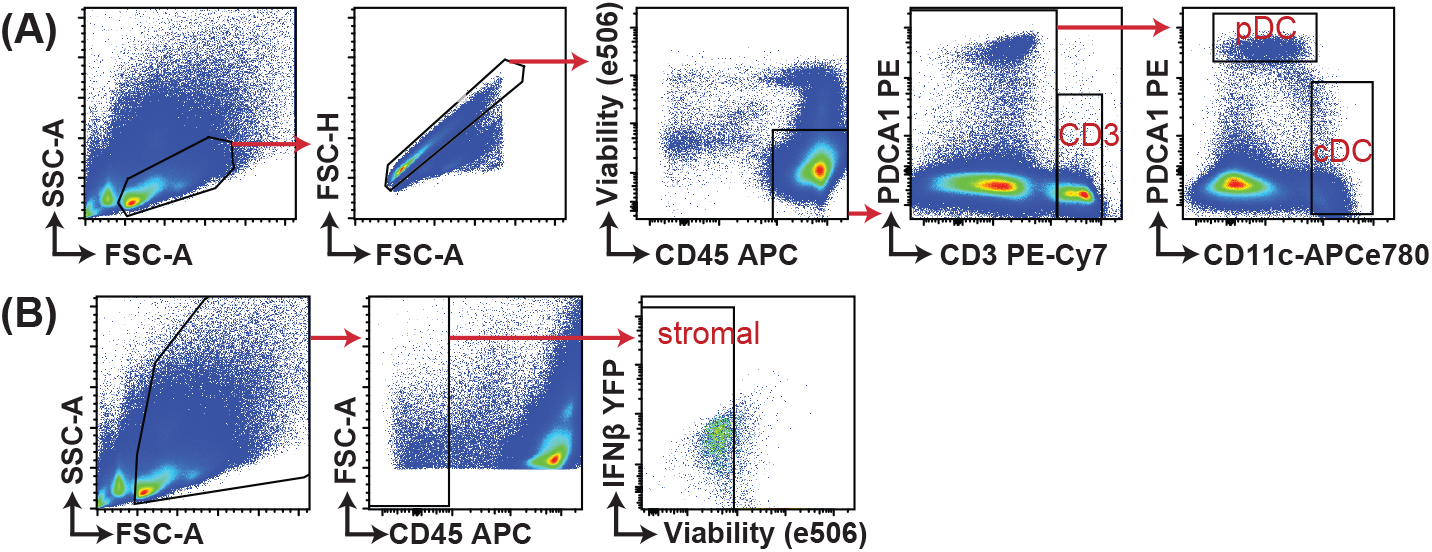
Gating strategy for analysis of IFNβ-YFP reporter mice. IFNβ-YFP reporter mice were backcrossed to MyD88-(MyD88 KO), STING-(STING GT) and double-deficient (DKO) mice. Animals were infected with 200,000 PFU WT1 MCMV and analyzed 48 hours post infection. Gating strategy for samples presented in Figure 4 are shown.

